# NF-kappa-B activation unveils the presence of inflammatory hotspots in human gut xenografts

**DOI:** 10.1101/2020.07.23.212621

**Authors:** Einat Nissim-Eliraz, Eilam Nir, Noga Marsiano, Simcha Yagel, Nahum Y. Shpigel

## Abstract

The single-epithelial cell layer of the gut mucosa serves as an essential barrier between the host and luminal microflora and plays a major role in innate immunity against invading pathogens. Nuclear factor kB (NF-κB), a central component of the cellular signaling machinery, regulates immune response and inflammation. NF-κB proteins are activated by signaling pathways downstream to microbial recognition receptors and cytokines receptors. Highly regulated NF-κB activity in intestinal epithelial cells (IEC) is essential for normal gut homeostasis; dysregulated activity has been linked to a number of disease states, including inflammatory bowel diseases (IBD) such as Crohn’s Disease (CD). Our aim was to visualize and quantify spatial and temporal dynamics of NF-κB activity in steady state and inflamed human gut. Lentivirus technology was used to transduce the IEC of human gut xenografts in SCID mice with a NF-κB luminescence reporter system. NF-κB signaling was visualized and quantified using low resolution, intravital imaging of the whole body and high resolution, immunofluorescence microscopic imaging of the tissues. We show that NF-κB is activated in select subset of IEC with low “leaky” NF-κB activity. These unique inflammatory epithelial cells are clustered in the gut into discrete hotspots of NF-κB activity that are visible in steady state and selectively activated by systemic LPS and human TNFα or luminal bacteria. The presence of inflammatory hotspots in the normal and inflamed gut might explain the patchy mucosal lesions characterizing CD and thus could have important implications for diagnosis and therapy.

## INTRODUCTION

Understanding mucosal immunity is critical for filling the knowledge gaps in the pathogenesis of inflammatory human gut disease such as Crohn’s Disease (CD). Most patients with Crohn’s disease have focal mucosal inflammation scattered throughout extensive portions of an otherwise normal bowel. It is assumed that gut inflammation occurs topographically at random and that the immune response potency is uniform within a region. The nuclear factor kappa B (NF-κB) family of transcription factors plays a central role in coordinating the expression of genes that control inflammation, immune responses, and a variety of other biological processes (Zhang et al., 2017). NF-κB plays a central role in IBD development and progression, and the level of activation of NF-κB correlates with the severity of intestinal inflammation (Atreya et al., 2008). Intestinal epithelial cells (IEC) are critical elements of gut homeostasis and function. Mikuda *et. al.* recently demonstrated that in IEC the intrinsic dual and seemingly opposing, anti- and pro-inflammatory activities of NF-κB are critical for normal gut homeostasis (Mikuda et al., 2020). Furthermore, these authors showed that dysregulated constitutive activity of NF-κB in IEC led to epithelial dysfunctions, barrier disruption and mucosal inflammation, all of which are known hallmarks of IBD.

Recent in vitro live-single-cell analysis of NF-κB activity revealed important molecular and mechanistic cues underlying the regulation of NF-κB (Patel et al., 2019;Son et al., 2019). These authors and others showed that activation of NF-κB in a seemingly uniform population of cells is highly heterogeneous and that cells are actually protected against harmful homogenous cellular activation (Adamson et al., 2016). Moreover, it was demonstrated that steady state cells with “leaky” NF-κB activity were high responders to various insults (Patel et al., 2019). Although understanding NF-κB activity in complex multicellular organs like the gut is a much more ambitious endeavor, some of these insights might help us to better understand the in vivo observations described here.

We have examined the activation of NF-κB in segments of human fetal gut transplanted and developing in a subcutaneous, sterile environment in SCID mice. The experimental model system we have used was first reported by Winter et al in 1991 (Winter et al., 1991) and refined in our laboratories for study of the human enteric nervous system (Nagy et al., 2018), human-specific pathogens (Golan et al., 2009;Golan et al., 2011) and inflammatory bowel diseases (IBD) (Canavan et al., 2016;Bruckner et al., 2019;Goldberg et al., 2019). Fetal gut was obtained after informed consent from pregnancy terminations performed legally at 12-18 weeks gestational age and transplanted subcutaneously in mature SCID mice. The human gut tissue vascularizes, expands and persists as a human gut implant, and can be experimentally manipulated over the course of the subsequent several months. While ectopic and not functional, these gut implants develop characteristic structures of the human gut with extensive vasculature and their structural features are highly similar to those of normal human gut, including mucosal villous epithelium, crypt structures, blood vessels and enteric nervous system.

We and others have previously shown that virtually all the cell types that are present in the normal human gut are also present in these xenografts (Savidge et al., 1995;Savidge et al., 2001;Golan et al., 2009;Golan et al., 2011;Nissim-Eliraz et al., 2017;Nagy et al., 2018;Bruckner et al., 2019). Moreover, the general architecture of the gut appears normal, and the tissue is well-vascularized by a human capillary system that anastomoses to the circulatory system of the murine host. In this, the experimental platform is similar to other examples of development of human tissues (lung, skin and liver) subcutaneously transplanted into SCID mice (Gaska and Ploss, 2015;Wahl et al., 2019). Furthermore, we have shown that many human innate and adaptive components of immune system, which have been shown to be already active in fetal gut at the time of transplantation (Rechavi et al., 2015;Li et al., 2018;Li et al., 2019;Schreurs et al., 2019;Stras et al., 2019), are present and active in the mature xenograft (Bruckner et al., 2019;Goldberg et al., 2019).

The model system allows the study of human immune activation in an animal model and is especially suited to the investigation of this response in mucosal epithelial of the gut. The intestinal epithelium is a single layer of cells with rapid turnover requiring constant tissue replenishment fueled by continuously dividing stem cells that reside at the bottom of crypts. These cells continuously proliferate and give rise to progenitor cells that differentiate to one of six different mature cell types and move upwards towards the villous, where they are shed into the intestinal lumen after 3-5 days (Gehart and Clevers, 2019). Henceforth, transduction of gut epithelium with a stable reporter gene requires infection of crypt stem cells and integration of the transgenes into their genome, all other transduced cells will undergo apoptosis and slough into the lumen.

HIV-derived lentiviral vectors (LVs) integrate into the target cell chromatin and are maintained as cells divide, a potential advantage for establishing long-term expression of reporter transgenes. We have developed LVs that achieve stable transgene expression in mucosal epithelial cells of human fetal gut xenografts. We monitored NF-κB dynamics by low resolution time-lapse bioluminescence imaging of human gut xenografts expressing a short-lived luciferase reporter gene controlled by NF-κB response elements.

## MATERIALS AND METHODS

### SCID mouse human intestinal xeno-transplant models

This model system was extensively described by us (Golan et al., 2009;Golan et al., 2011;Canavan et al., 2016;Nissim-Eliraz et al., 2017;Nagy et al., 2018;Bruckner et al., 2019;Goldberg et al., 2019). C.B-17/IcrHsd-*Prkdc*^*scid*^ (abbreviated as SCID) mice were purchased from Harlan Biotech Israel (Rehovot, Israel). All mice were housed in a pathogen-free facility, in individually ventilated cages (IVC), given autoclaved food and water. All animal use was in accordance with the guidelines and approval of the Animal Care and Use Committee (IACUC) of the Hebrew University of Jerusalem. IRB and IACUC approvals were obtained prospectively (Ethics Committee for Animal Experimentation, Hebrew University of Jerusalem; MD-11-12692-4 and the Helsinki Committee of the Hadassah University Hospital; 81-23/04/04). Women undergoing legal terminations of pregnancy gave written, informed consent for use of fetal tissue in this study.

Human fetal small bowel 12-18 weeks gestational age was implanted subcutaneously on the dorsum of the mouse as described previously (Golan et al., 2009;Golan et al., 2011). All surgical procedures were performed in an aseptic working environment in a laminar flow HEPA-filtered hood with isoflurane inhalation anesthesia (1.5 to 2% v/v isofluorane in O_2_). Before surgery, carprofen (5 mg/kg, Rimadyl, Pfizer Animal health) was administered subcutaneously. The surgical area was shaved and depilated (Nair hair removal cream) and the skin was scrubbed and disinfected with betadine and 70% (v/v) ethanol. After surgery the mice were provided with enrofloxacin-medicated water (Bayer Animal HealthCare AG) for 7 days and were closely monitored once a day for behavior, reactivity, appearance and defecation. Grafts developed in situ for 12-16 weeks prior to manipulation.

### Activation of inflammation

We used LPS (from *E. coli* serotype O55:B5, Sigma, Rehovot, Israel) or human TNFα (Peprotech) by intraperitoneal injection or intraluminal bacterial infection to induce inflammatory response in the implant in SCID mice with mature human gut xenograft. Transplanted mice were subjected to systemic LPS (0.1, 1 or 10 mg/kg) or human TNFα (1 mg/kg), well-established models of experimental sepsis and inflammatory gut disease (Vereecke et al., 2010;Guma et al., 2011;Van Hauwermeiren et al., 2013;Williams et al., 2013;Newton et al., 2016). Enteropathogenic *E. coli* (EPEC) bacteria, serotype O127:H6 strain E2348/69 (Nataro & Kaper, 1998), were used in this study. Bacteria were grown in Luria-Bertani (LB) broth at 27°C. For in vivo challenge studies, cultures were diluted to 10^7^ CFUs in 100 µl culture medium and injected into the lumen of the human gut xenografts.

### Plasmids and Recombinant Viruses

To create lentiviral NF-κB reporter vectors, two target plasmids (transfer vectors) were constructed to express a low resolution luminescence reporter and a high resolution fluorescence reporter. The luminescence reporter was constructed of destabilized firefly luciferase (pGL4.24-luc2P, Promega) (Li et al., 1998;Suter et al., 2011) open reading frame controlled by a DNA cassette containing five tandem repeats of the NF-κB transcriptional response element (pLNT-minP-5κB-luc2P) (Brignall et al., 2017). The fluorescence reporter was constructed of human p65 N-terminal fusion with tagRFP reporter (pLNT-UbC-tagRFP-p65). The lentiviruses were produced from HEK293FT cells (ThermoFisher scientific) through co-transfecting the target plasmid pLNT and the packaging vectors pMDLg-RRE, pRSV-REV and pCMV-VSVG as previously described (Bagnall et al., 2015). Co-transfection using VSVG plasmid encoding for envelope G glycoprotein from VSV, produced pseudotyped retrovirus which depends on the expression of Low density lipoprotein receptor (LDLR) on plasma membrane for the infection of target cells (Finkelshtein et al., 2013;Amirache et al., 2014;Girard-Gagnepain et al., 2014). Lentiviruses were purified by ultracentrifugation and then quantitated.

### In vitro lentivirus target validation (Supplementary Figures S1 and S2)

The lentivirus target and reporter system were validated in mouse and human tissue cultured cell lines (Salamon et al., 2020). Mouse RAW 264.7 and human THP-1 macrophage-like cell lines, mouse mammary epithelial line EPH4 and human embryonic cell line HEK293FT were infected with the lentiviral NF-κB reporter vector. RAW 264.7 and THP-1 cells were cultured and activated by LPS as previously reported (Mintz et al., 2013).

EPH4 and HEK293T cells were cultured in Dulbecco’s modified Eagle medium (DMEM) supplemented with 10% fetal calf serum (FCS) (Biological Industries) and 100 units/ml penicillin, 0.1 mg/ml streptomycin (Pen-Strep; Biological Industries), 1% L-Glutamine (BI) and 1% HEPES buffer (BI) at 37°C with 5% CO_2_. EPH4 and HEK293T cells were infected with the following strains of Enteropathogenic *E. coli* (EPEC); EPEC wild-type E2348/69, E2348/69Δ*escV*::*kan*, E2348/69ΔIE6::*cm* ΔPP4::*kan* and E2348/69ΔnleBCDE (kindly donated by I. Rosenshine, Hebrew University) (Baruch et al., 2011). Bacteria were grown in Luria-Bertani (LB) broth at 27°C. For in vitro challenge studies, bacterial cultures were diluted in cell culture medium to multiplicity of infection (MOI) of 1. Cells were plated at equal density and luminescence signals were quantified in the presence of 150 µg/ml D-luciferin (GoldBio) by imaging using cooled CCD optical macroscopic imaging system (IVIS Lumina Series III, PerkinElmer Inc., MA USA) and SpectraMax i3x multiple detection microplate reader (Molecular Devices, CA USA).

Fluorescence *in situ* hybridization (FISH) was performed using Stellaris RNA FISH (LGC Biosearch Technologies, Petaluma, CA) as previously described (Raj et al., 2008). A set of Quasar 670-labeled oligonucleotides probes were designed to selectively bind to luc2P transcripts. EPH4 cells transduced with pLNV-minP-5kB-luc2P were grown on 12 mm glass coverslips in 24-well plates at a density of 3×10^5^ cells/well. The next day, cells were treated with 10 µg/ml *Mycoplasma bovis* lipoproteins for 2hr at 37°C. *Mycoplasma bovis* strain 161791 (Lysnyansky et al., 2016) lipoproteins were prepared using Triton X-114 phase fractionation method as previously described (Elkind et al., 2012). Hybridizations were done according to manufacturer protocol for adherent cells, using commercial hybridization buffer and wash buffers, 4,6-diamidino-2-phenylindole (DAPI) dye for nuclear staining was added during the washes. Images were taken with epifluorescence microscopy system (Axio Imager M1, Zeiss, Germany).

### Lentivirus injection

Intra-xenograft viral injections were performed in mature gut xenografts 12-16 weeks after transplant. Mice were prepared and anesthetized as described above, using 27 gauge beveled needle the virus was injected directly into the wall of the gut xenograft through the skin above the transplant. Each xenograft received 10×15 µl injections of lentivirus applied along the long axis of the dorsal wall.

### Real-time PCR for analyses of Luciferase gene copy number in human gut xenografts

Total genomic DNA was extracted from the focus as well as quiescent regions of the transduced xenograft using DNeasy Blood and Tissue kit (Qiagen, Hilden, Germany), according to the manufacturer’s instructions. To estimate the copy number of the Luc2p gene in the samples, 100 ng genomic DNA was added to the FAST qPCR Universal Master Mix (Kappa Biosystems, Boston, MA, USA) with Luc2p specific primers 5’-TGCAAAAGATCCTCAACGTG-3’ (forward) and 5’-AATGGGAAGTCACGAAGGTG-3’(reverse) and quantitative real-time RT–PCR was conducted on a StepOne Plus PCR instrument (Applied Biosystems). The NF-κB-luciferase (Luc2p) reporter construct plasmid used for standard curve generation was diluted with genomic DNA from untransduced xenograft to control for any inhibitory effect of genomic DNA on PCR. The number of vector DNA molecules in transduced cells was calculated by comparing threshold cycle (Ct) values of samples to that of the plasmid standard curve. For determining the final DNA titer of vectors, the total number of vector DNA molecules in transduced xenografts was normalized to the number of genomes determined by the PTBP2 gene previously described (Salguero et al., 2014).

### Bioluminescence and fluorescence Imaging

For in vivo imaging, mice were anesthetized and given D-luciferin by IP (150 mg/kg) and whole body imaging was performed using IVIS Lumina Series III (PerkinElmer Inc., MA USA) (Prescher and Contag, 2010). As previously described (Evans et al., 2014), all images were analyzed using Living Image software (version 4.4, Caliper LifeSciences (MA, USA) and the optical signal intensity was expressed as photon flux in units of photons/s/cm^2^/steradian. Each image was displayed as a false-color photon-count image superimposed on a grayscale anatomic image. To quantify the measured light, regions of interest (ROI) were defined over the subcutaneous human gut xenografts. All values were examined from an equal ROI.

### Histochemical analysis

As we have previously described (Nissim-Eliraz et al., 2017), mice were killed at the indicated time points after challenge and xenograft tissues were trisected for histology and fluorescence staining. Samples for histological analysis were fixed in neutral buffered 4% paraformaldehyde (PFA), paraffin embedded, and sections were cut at a thickness of 5 µm and stained with hematoxylin and eosin (H&E). Fresh xenograft tissue for fluorescence staining was fixed in 2.5% PFA overnight at room temperature, incubated with 15% (w/v) sucrose for 12 hours at 4°C and frozen in Tissue-Tek® (EMS, Hatfield, PA) embedding medium. Serial 10 µm cryosections were stained with phalloidin (Sigma) and 4’, 6’-diamidino-2-phenylindole (DAPI) (Sigma). For immunofluorescence staining primary antibodies were anti-luciferase antibody (ab21176 Abcam, Cambridge, UK) and alexa fluor 594 Donkey anti rabbit (A21207 Invitrogen, Carlsbad, CA) were used as secondary antibodies. Sections were mounted with VectaShield (Vector Laboratories,Burlingame, CA) and imaged with an Axio Imager M1 upright light microscope (Zeiss, Germany) coupled to a MR3 CCD camera system (ZEN 2012).

### Quantitative RT-PCR for analysis of LDLR and PCSK9

Low density lipoprotein receptor (LDLR) and proprotein convertase subtilisin/kexin type 9 (PCSK9) genes expression in human gut xenografts was analyzed using QPCR as previously described (Nissim-Eliraz et al., 2017;Bruckner et al., 2019). Briefly, total RNA was isolated from xenograft tissue using the GeneElute Mammalian Total RNA Miniprep Kit (Sigma, Rehovot, Israel) combined with on-Column DNase I Digestion Set (Sigma). Reverse transcription was performed using qScript cDNA Synthesis Kit (Quanta biosciences, Gaithersberg, MD, USA) and cDNA was used for subsequent real-time PCR reactions. Quantitative real-time RT–PCR was conducted on a StepOne Plus PCR instrument (Applied Biosystems) using the FAST qPCR Universal Master Mix (Kappa Biosystems, Boston, MA, USA). Primer pairs were: LDLR FORW - 5’ GTCTTGGCACTGGAACTCGT, and LDLR REV - 5’CTGGAAATTGCGCTGGAC; PCSK9 FORW - 5’AGGGGAGGACATCATTGGTG, and PCSK9 REV - 5’CAGGTTGGGGGTCAGTACC. All reactions were performed in triplicate, and the gene expression levels for each amplicon were calculated using the 2^-ΔΔCT^ threshold cycle (CT) method (Livak and Schmittgen, 2001), and the levels were normalized against those for human β_2_-microglobulin (B2M) mRNA. Melting curve analysis was performed with each primer set to confirm amplification of a single product, and all amplicons were sequenced to ensure reaction specificity (data not shown).

## RESULTS

### LDLR expression in human gut xenografts

Our initial attempts to efficiently infect and transduce cells in human fetal gut or fully developed xenografts by injection of lentivirus into the lumen or directly into the gut wall failed. Transduction of human fetal gut cells prior to transplantation was a rare event and difficult to locate using fluorescence microscopic imaging of cryosections. Nevertheless, these rare events clearly demonstrated that the heritable genetic marker p65-RFP was integrated into crypt stem cells via lentiviral vector infection, and that all other cells if at all transduced by the lentivirus vector were normally turned over and lost in the course of xenograft development over a 12 week period. The transduced stem cells in which the p65-RFP transgene have been integrated generated progeny that produce the p65-RFP protein. These cells extend from the crypt base to the villus tip and can be visualized as ribbons of red fluorescing cells (Supplementary Figure S3). In attempt to improve the efficiency of lentivirus virus infection and cell transduction we analysed the expression of LDLR expression in human fetal gut and fully developed xenografts. LDLR was previously reported to serve as the major entry port of VSV-G–pseudotyped lentiviral vectors in human and mouse cells (Finkelshtein et al., 2013;Amirache et al., 2014;Girard-Gagnepain et al., 2014;Kohn and Hollis, 2014). Furthermore, it was also demonstrate that LDLR is present in gut mucosal epithelial cells and localized to the basolateral membrane (Fong et al., 1989;Pathak et al., 1990;Levy et al., 2013;Meoli et al., 2018). Thus, we have used QPCR to analyze the expression of LDLR in the mature human gut xenograft. Surprisingly, the expression of LDLR in mature human gut xenografts was very low in steady state (Figure 1A-B). This dearth of LDLR expression in mature human gut xenografts and the lack of expression on the apical membrane of the enterocytes probably explain our failure to infect and transduce these cells.

**Figure 1.**
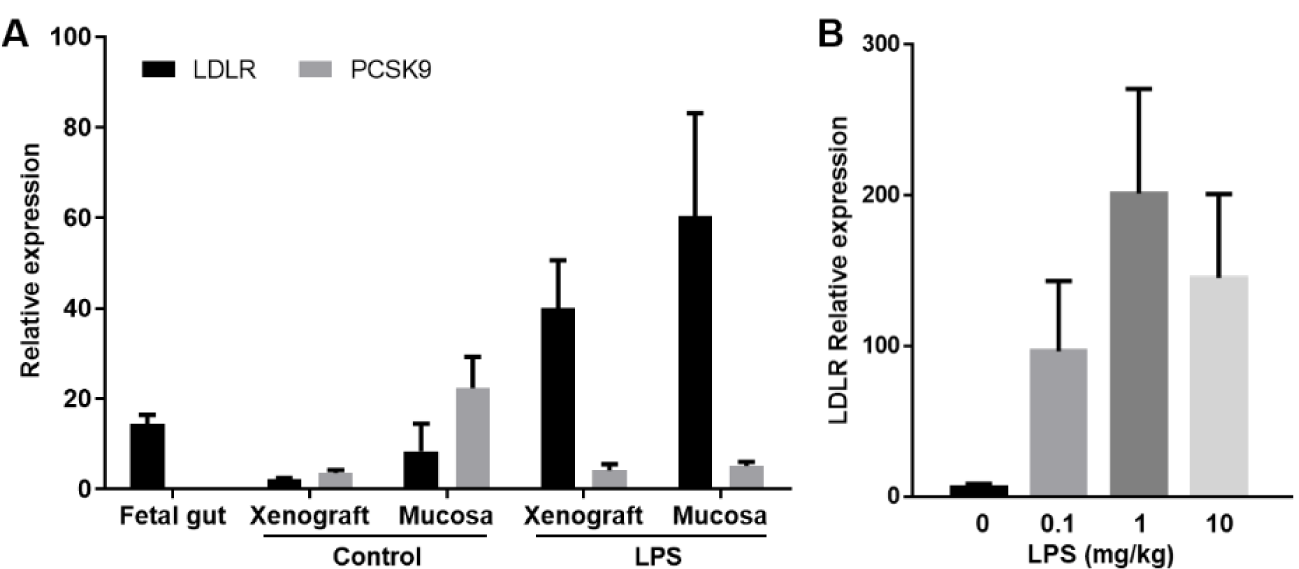
LPS treatment increase LDLR and suppress PCSK9 genes expression in human gut xenografts. Human fetal gut was transplanted into SCID mouse and allowed to develop over 12 weeks. Mice with fully developed gut xenografts were systemically treated with 10 mg/kg LPS by intraperitoneal injection (A). Dose response experiments (0.1-10 mg/kg LPS) demonstrated the utility of lower LPS dose (0.1 mg/kg) to increase the expression of LDLR in human gut xenografts (B). Normal fetal gut before transplantation, and normal and LPS-treated whole xenografts and mucosal scrapings were harvested for RNA extraction and QPCR analysis 4 hours after LPS treatment or PBS as control treatments. Graphs show mean ± SEM of relative expression.

In vitro and in vivo studies demonstrated increased gene expression and protein production of LDLR by LPS and inflammatory cytokines in many cell types (Chen et al., 2007;Li et al., 2013). Furthermore, a recent study established the significant role of LDLR as LPS scavenger in sepsis animal models and clinical situations (Walley et al., 2014). More importantly, this study also demonstrated the inhibitory effect of proprotein convertase subtilisin/kexin type 9 (PCSK9) on LPS scavenging and clearance in sepsis. PCSK9, a serum protein which is also increased by LPS and in sepsis, binds to LDLR stimulates its internalization, promotes LDLR lysosomal degradation, and prevents recycling of LDLR to the cell surface. Thus we hypothesize that LPS might increase cells surface density of LDLR in the human gut xenografts, albeit, this effect might be negated by increased human and mouse PCSK9 which are cross reactive. To test this hypothesis, we administered LPS intraperitoneally (IP) to SCID mice with mature human gut xenografts to measure the expression of LDLR and PCSK9. We have found that systemic administration of LPS to the host mouse resulted in significant increase of LDLR expression in the mature human gut xenografts and decreased expression of PCSK9 expression (Figure 1A). Consequently, we established our working protocol where lentivirus carrying luciferase gene under the control of 5 copies of the NF-κB response element (NF-κB/RE-luciferase reporter; Supplementary Information, Figure S2A) was injected into the wall of fully-developed human gut xenografts 4-6 hours after administration of systemic low dose of LPS (0.1 mg/kg; Fig. 1B) to the host mouse.

### Intravital imaging of NF-κB activity in the human gut

Human fetal gut segments were transplanted subcutaneously in SCID mice and allowed to develop over 12-16 weeks thereafter. Expression of LDLR was activated in fully developed transplants by treating the host mouse with intraperitoneal (IP) injection of 0.1 mg/kg LPS and Lentivirus (NF-κB/RE-luciferase reporter) was injected into the human gut wall 4-6 hours thereafter. Mice were allowed to recover and 2-3 weeks thereafter were subjected to intravital whole body bioluminescence imaging before and after systemic IP injection of LPS (Figure 2) or human TNFα (Figure 3) and intraluminal challenge with wildtype (WT) or T3SS-defective mutant enteropathogenic *E. coli* (EPEC) bacteria (Figure 4). We have previously demonstrated activation of acute inflammation in human gut xenografts following systemic LPS (Bruckner et al., 2019) or intraluminal EPEC infection (Nissim-Eliraz et al., 2017). While the tacit assumption is that gut inflammation is homogeneous, we surprisingly observed one to two foci of luminescence activity in all xenografts following challenge with similar time course for LPS and TNFα and somewhat longer activity following luminal bacterial challenge. In some xenografts focal activity was detected even before challenge (Figure 3A and Figure 4B) and these steady state foci were further activated following challenge.

**Figure 2.**
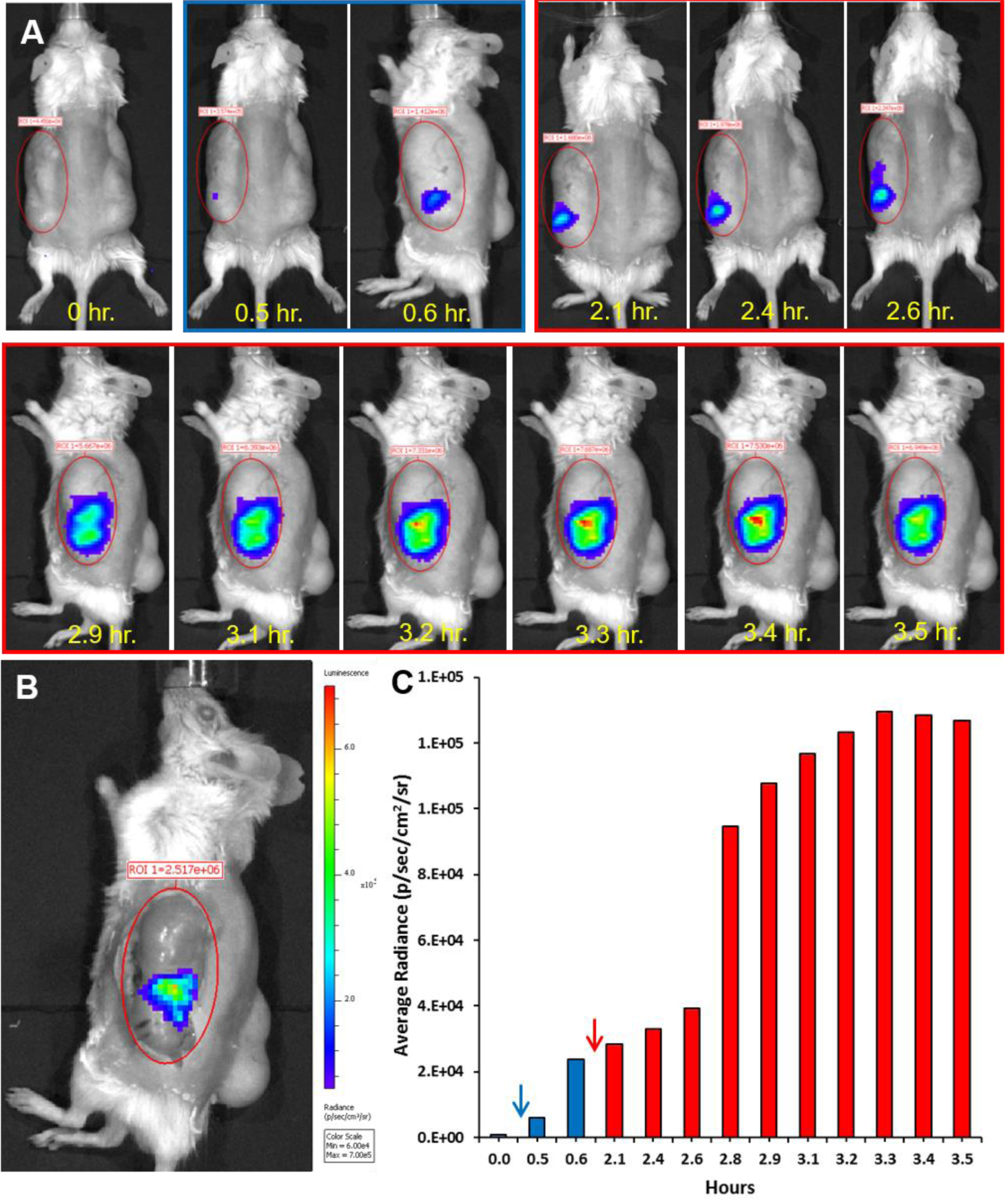
Spatial and temporal activity of NF-kappa-B in steady state and inflamed human gut activated by systemic LPS. Human fetal gut was transplanted into SCID mouse and allowed to develop over 12 weeks. Using lentivirus vector, mature human gut xenografts were transduced with luciferase gene under the control of 5 copies of the NF-κB response element. Two weeks thereafter, luminescence intravital imaging was performed using cooled CCD optical macroscopic imaging system (IVIS Lumina Series III, PerkinElmer Inc., MA USA) and the time course of the luminescence activity before activation (blue frame in A) and after activation (red frames in A) are presented. To quantify the measured light, images were analyzed using Living Image software (version 4.4, Caliper LifeSciences, MA, USA), equal regions of interest (ROI) were defined over the subcutaneous human gut xenografts (red ellipses in A-B) and the optical signal intensity was expressed as photon flux in units of photons/s/cm2/steradian (color scale in B and bar graph in C). While no luminescence signal was observed before systemic injection of D-luciferin (time point 0 hr. in upper left panel in A and bar graph in C), steady state focal activity was visible (blue frame in A and blue bars in C) after luciferin injection (blue arrow in C). The focal activity of NF-κB was discretely activated (red frames in A and red bars in C) following systemic injection of LPS (1 mg/kg) (red arrow in C). To further verify the origin of luminescence signal, the skin above the xenograft was removed and the exposed xenograft was imaged in the live mouse (B). Representative results of one experiment out of five similar experiments.

**Figure 3.**
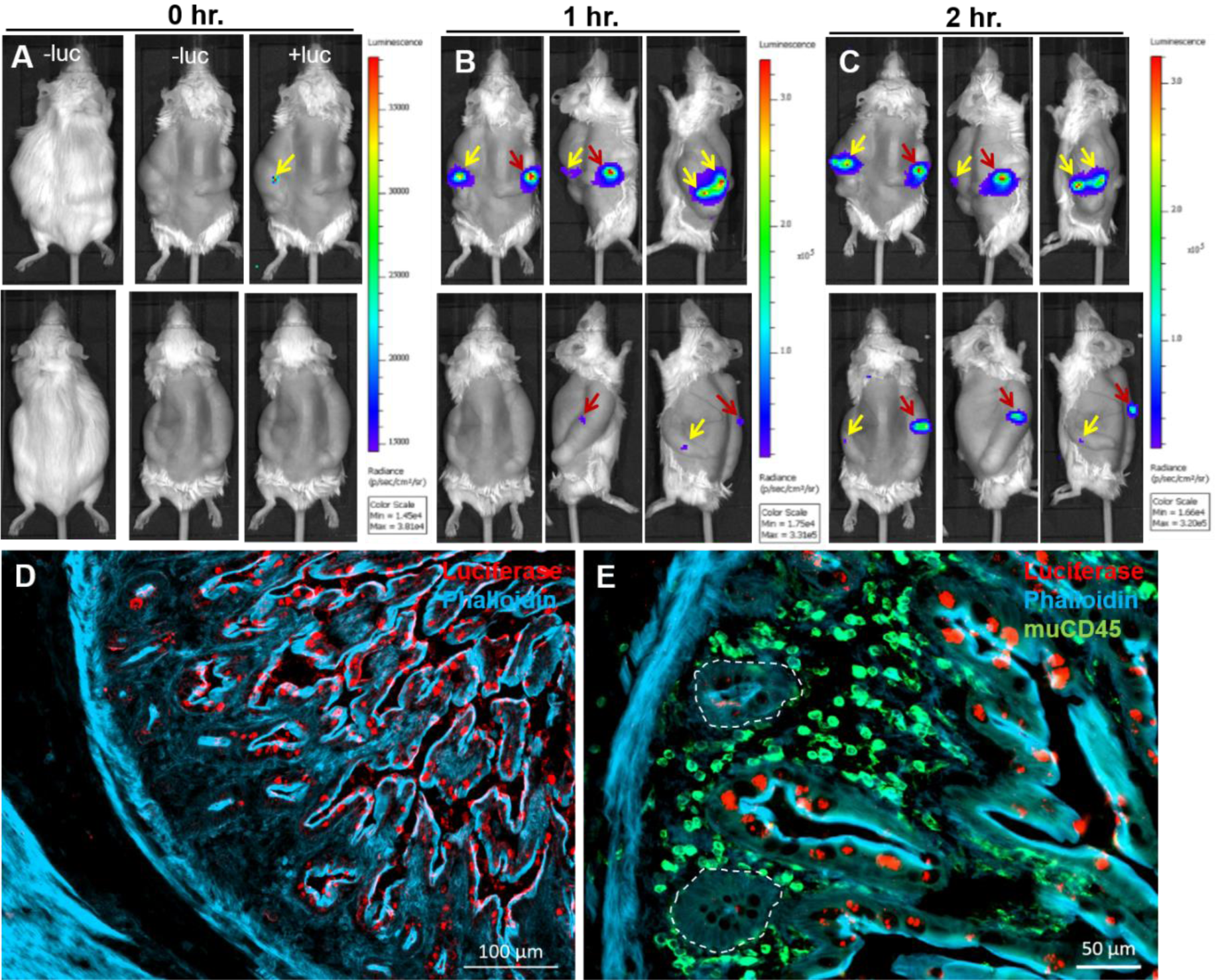
Spatial and temporal activity of NF-κB in steady state and inflamed human gut activated by systemic human TNFα. Human fetal gut was transplanted into two SCID mice and allowed to develop over 12 weeks. The reporter system and intravital imaging were performed as described above (see Figure 1). Time course of the luminescence activity before activation (A) and 1 (B) and 2 (C) hours after activation following systemic injection of human TNFα are presented. While no luminescence signal was observed before systemic injection of D-luciferin (-luc in A), steady state focal activity was visible in one transplant (yellow arrow in A) after luciferin injection. The focal activity of NF-κB was discretely activated following systemic injection of TNFα (1 mg/kg) (yellow and red arrows in B-C). The expression of luciferase in the human gut transplant 2 hours after activation with TNFα was imaged using immunostaining and fluorescence microscopy (D-E). Representative images of fluorescence staining of cryosections using phalloidin (D-E, blue), anti luciferase antibodies (D-E, red) and anti murine CD45 antibodies (green in E). Luciferase expression is limited to mucosal villous epithelial cells (crypts are demarcated by broken white line E). Scale bars, 100 µm (D) and 50 µm (E). Representative results of one experiment out of five similar experiments.

**Figure 4.**
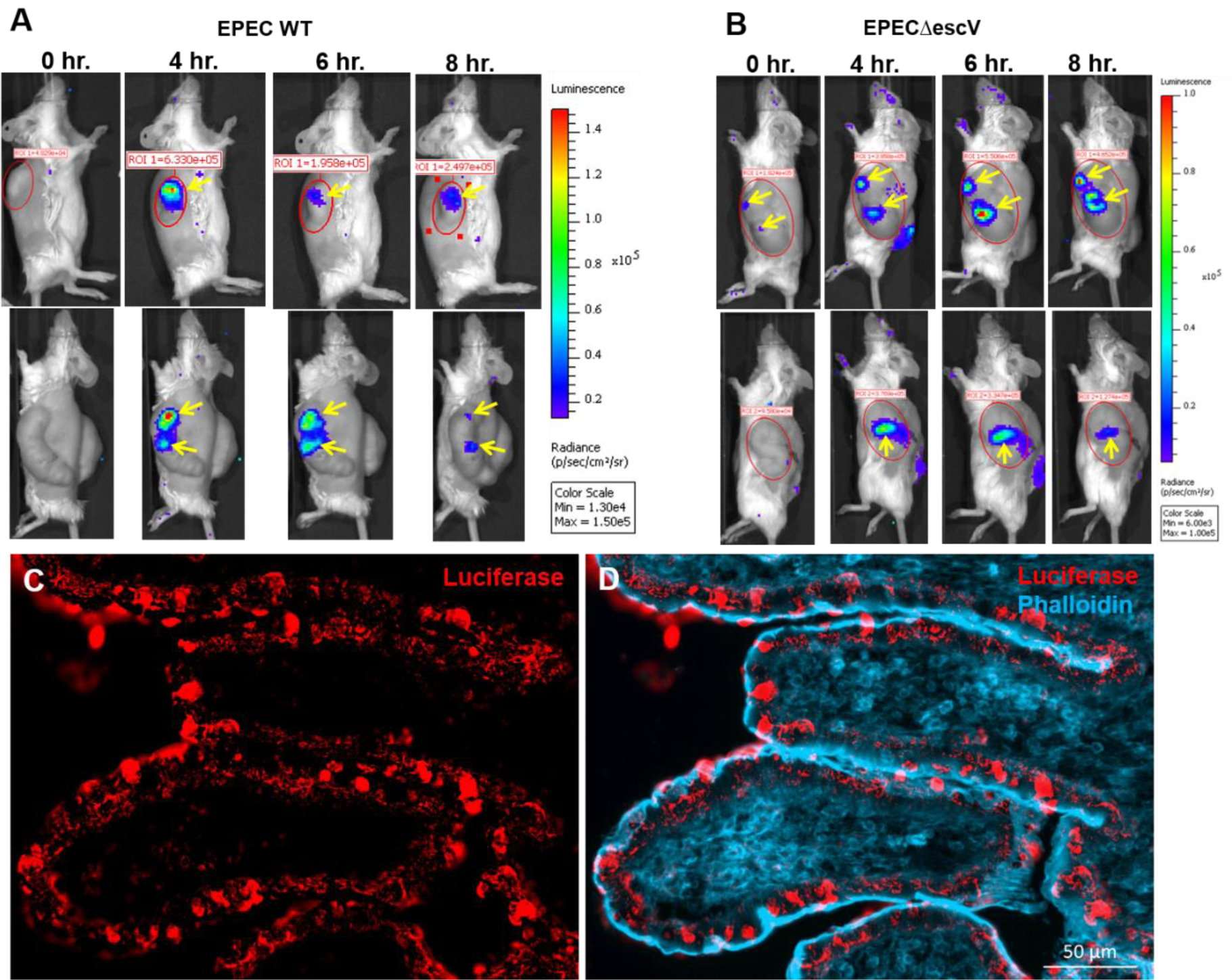
Spatial and temporal activity of NF-κB in steady state and inflamed human gut activated by luminal *E. coli* bacteria. Human fetal gut was transplanted into two SCID mice and allowed to develop over 12 weeks. The reporter system and intravital imaging were performed as described above (see Figure 1). Time course of the luminescence activity before activation (0 hr. in A-B) and 4-8 hours (A-B) after activation following intraluminal infusion of Enteropathogenic E. coli O127 wildtype (WT) strain (A) and the T3SS-defective isogenic Δ*escV* mutant strain (B). Focal activity in steady state was visible in one transplant (yellow arrow in B) following luciferin injection. Luminescence imaging demonstrating focal activity of NF-κB discretely activated 4-8 hours following intraluminal infection with WT or mutant bacterial strains (yellow arrows in A-B). The expression of luciferase human gut transplant 8 hours after infection with WT bacteria was imaged using immunostaining and fluorescence microscopy (C-D). Representative images of fluorescence staining of cryosections using anti-luciferase antibodies (red in C-D) and phalloidin (blue in D). Luciferase expression is limited to mucosal villous epithelial cells. Scale bars, 50 µm (C-D). Representative results of one experiment out of five similar experiments.

Activated xenografts were harvested and luciferase expression was further analyzed using immunofluorescence microscopy (Figure 3D-E and Figure 4C-D). Luciferase was only detected in foci of luminescence activity and was limited to mucosal epithelial cells of the human gut.

The unexpected observation of focal NF-κB activity in steady state and inflamed human gut might represent foci of higher lentivirus infection and transduction efficiency. To address this possibility we established a real-time quantitative PCR system for analyses of luciferase gene copy number in the human gut xenografts (Figure 5). Our analysis clearly showed that foci of NF-κB activity were not associated with high copy number of luminescence reporter. Viral infection and reporter gene transduction were homogeneously distributed along the human gut xenografts thus verifying that NF-κB activity in mucosal epithelial cells of steady state and inflamed human gut is focal.

**Figure 5.**
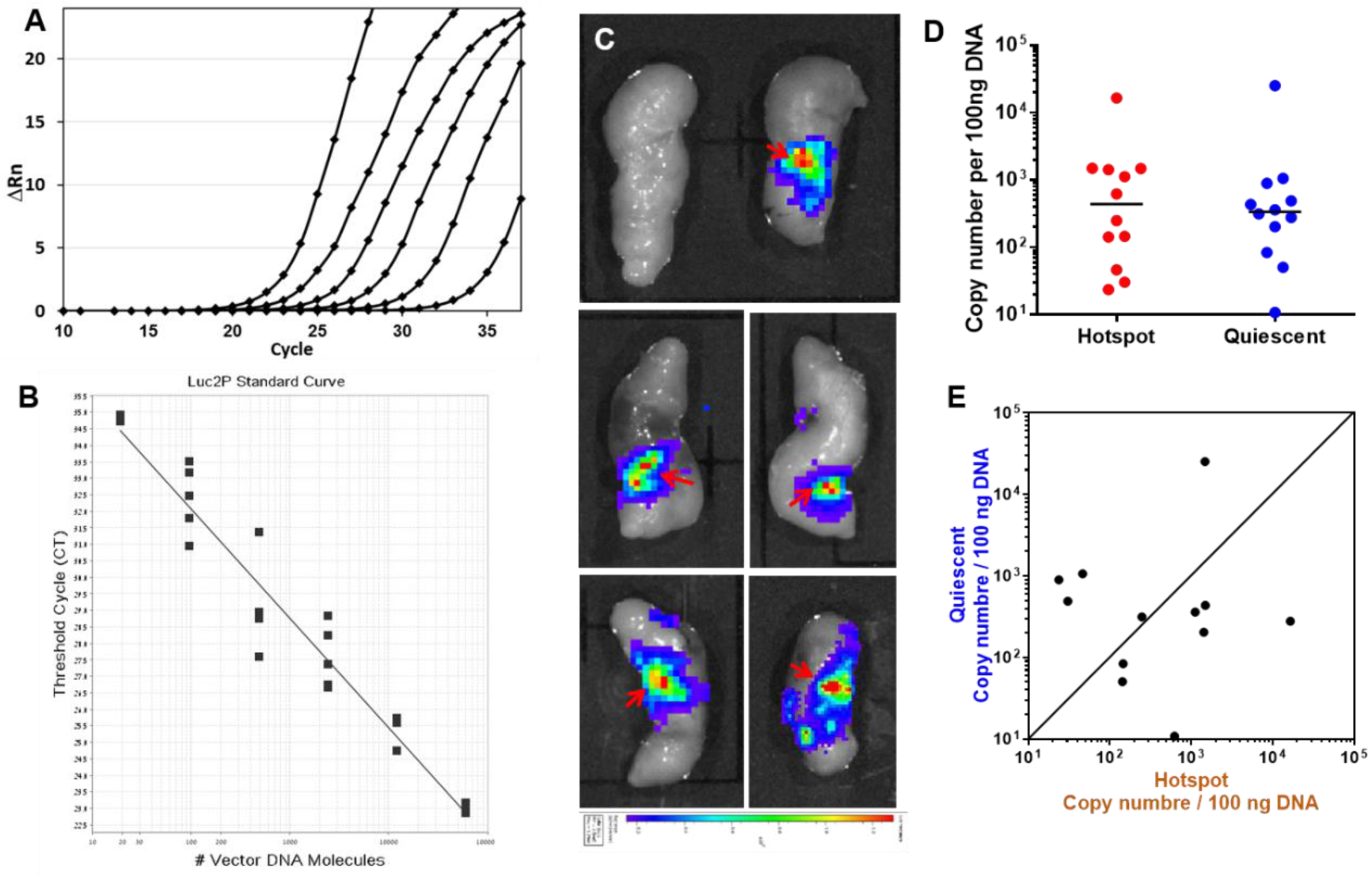
Foci of NF-κB activity are not associated with high copy number of luminescence reporter. Real-time PCR was used for analyses of Luciferase gene copy number in human gut xenografts. Amplification of the NFκB-Luciferase plasmid (A) and derived standard curve (B). The plasmid was serially diluted 1:5 starting from 6×10^4^ vector DNA molecules and was amplified using the Luc2P primers. Plasmid DNA was diluted with genomic DNA to generate the standard curve (B) as described in Materials and methods. The amplification plot shows the change in fluorescence (Rn) as a function of the PCR cycle (A). Gut transplants were imaged ex-vivo for luminescence activity as described above (see Figure 2) to enable accurate sampling of hotspots (red arrows in C) and quiescent regions for extraction of genomic DNA from transduced and non-transduced transplants (top left panel in C) as negative controls. Copy number of reporter genes was not consistently higher in the luminescence foci compared with quiescent regions of NF-κB activity in transduced transplants (D-E). Each data point represents sampled transplant in luminescence focus (red hotspots in D) or quiescent area (blue in D). Paring of hotspot and quiescent copy number in individual transplants is presented in E.

## DISCUSSION

NF-κB, a rapidly inducible transcription factor, is the master regulator of inflammation in all body systems and implicated in the pathogenesis of disease conditions such as human IBD (Zhang et al., 2017). Previous animal studies demonstrated that impairment of NF-κB signaling in intestinal epithelial cells resulted in inflammatory diseases. To this end, better understanding the temporal and spatial dynamics of NF-κB activity is important and these were previously analyzed in various animal models. Using transgenic mice that express luciferase under the control of NF-κB, intravital activation of NF-κB was visualized in skin, lungs, spleen, Peyer’s patches, and the wall of the small intestine at 4-5 hours following systemic treatment with TNFα, IL1α and LPS (Carlsen et al., 2002). This system enabled low resolution, intravital, spatial and temporal analysis of NF-κB activity in superficial structures like skin and joints or ex-vivo snap shots of isolated organs. However, the strong signaling emanating from lymphoid tissues, Peyer’s patches, and gut parenchymal cells precluded specific analysis of NF-κB activity in gut mucosal epithelial layer. Alternatively, using transgenic mice that express enhanced green fluorescence protein (EGFP) under the control of NF-κB, enabled high resolution microscopic analysis on sections of the gut (Magness et al., 2004). These mice were systemically treated with LPS and gut sections were microscopically analyzed for EGFP fluorescence 4 and 24 hours thereafter. These authors suggested regional specificity in NF-κB responsiveness in the gut which was mostly attributed to immune cells in the lamina propria of the duodenum and proximal jejunum. Interestingly, gut epithelial cells were only sparsely activated although small foci of activated epithelial cells are visible in the published images. These observations were further supported by studies in NF-κB/EGFP transgenic zebra fish demonstrating NF-κB activation in subpopulations of intestinal epithelial cells by luminal bacteria (Kanther et al., 2011).

Although technically challenging, using low resolution bioluminescence imaging and high resolution fluorescence microscopy, the above described experimental platforms can be used for intravital spatial and temporal analysis of NF-κB activity in steady state and inflamed mouse gut. Furthermore, this can be achieved in the same animal using transgenic mice co-expressing luciferase and EGFP regulated by a bidirectional NF-κB response element as previously reported (Kielland et al., 2012). Moreover, cell-specific (e.g. gut epithelial cells) co-expression of bioluminescence and fluorescence reporters can be achieved in transgenic mice using the Cre-LoxP technology.

Here, we have used the xenograft model system to study the dynamics of NF-κB in steady state and inflamed human gut. This experimental platform also enabled us to circumvent some of the above described limitations. We show here that genetic manipulation of human gut cells can be achieved using lentivirus technology and that the ectopic subcutaneous location render the transplants highly accessible for accurate viral infection, bacterial intraluminal challenge and intravital imaging. As VSV-G lentivirus infection is limited to cells expressing LDLR and is inefficient in lymphoid (Amirache et al., 2014) and myeloid immune cells (Milani et al., 2019), we have found that infection and transduction were limited to the mucosal epithelial cells in the human gut xenografts. Henceforth, this lentivirus technology enabled us to transduce mucosal epithelial cells of human gut transplants with a genetic construct containing five NF-κB binding sites coupled to the gene encoding firefly luciferase. We have used this experimental platform to study the temporal and spatial dynamics of NF-κB in steady state and following systemic activation with LPS or TNFα, or intraluminal challenge with a human-specific Enteropathogenic *E. coli*. Surprisingly, intravital bioluminescence imaging of NF-κB activity in gut epithelial cells exposed “hotspots” representing clusters of NF-κB-activated cells. We suspected that these foci of luciferase expression might be related to inhomogeneous viral injection and transduction rates, however, luciferase gene copy number analysis was not supportive for this plausible explanation.

We have also conducted a more detailed analysis of hotspots and quiescent gut tissues using anti luciferase immunofluorescence microscopy. This extensive and exhaustive analysis of the tissues revealed three important observations; (1) luciferase expression was not detected in the quiescent gut tissues, (2) luciferase expression could only be detected in human gut epithelial cells and was absent from other cell types in the xenografts, including from CD45-positive immune cells, (3) luciferase expression was highly variable among epithelial cells within individual hotspots. To this end, we have noticed that Karrasch et al (Karrasch et al., 2007) presented confocal images of colonic sections depicting similar transient and variable NF-κB activity in intestinal epithelial cells from NF-κB/EGFP transgenic mice following bacterial colonization. Furthermore, these results are also consistent with previous animal studies demonstrating constitutive NF-κB activity in subsets of normal gut epithelial cells which are further activated by bacteria and inflammatory mediators (Kanther et al., 2011). Further work is required to better characterize NF-κB activity in distinct cell types of the intestinal epithelial interface.

Although our results cannot be directly compared with in vitro studies of NF-κB activity in other cell types, the hotspots of NF-κB activity observed in steady state xenografts might represent the highly responsive “leaky” cells reported by Patel et al. (Patel et al., 2019). It remains an open question as to what may leads to this variation in NF-κB activity in steady state IEC, one intriguing possibility is that epigenetic variance in genes encoding for NF-κB network components enforce this variability. Indeed, epigenetic mechanisms are known to shape gene expression in ISC (Hu et al., 2019), the progenitors of all IEC, and were linked to susceptibility IBD (Howell et al., 2018). It still remain to understand if hotspots of NF-κB activity are normal feature of the human gut or we can also speculate that epigenetic changes affecting the developing human fetal gut as subcutaneous xenografts in SCID mice might lead to the development of clusters of “leaky” epithelial cells hyper responsive to general insults. Either way both of these notions are plausible and further studies are required to elucidate the underlying cellular and molecular mechanisms involved and to understand their relevance to clinical situations in IBD patients.

## ACKNOWLEDGEMENTS

The authors acknowledge the technical assistance in transplantation of human fetal gut from Dr. Irit Shoval.

The authors acknowledge the scientific and technical assistance in the construction and use of the lentivirus technologies from Drs. James Bagnall and Pawel Paszek of the Faculty of Biology, Medicine and Health, University of Manchester, Manchester, United Kingdom.

## AUTHOR CONTRIBUTIONS

N.Y.S designed the experiments, E.N.E, E.N and N.M performed the experiments, S.Y. provided human tissues, N.Y.S analyzed the data and wrote the manuscript with inputs from all authors.

## Figure legends

**Supplementary Figure S1.**
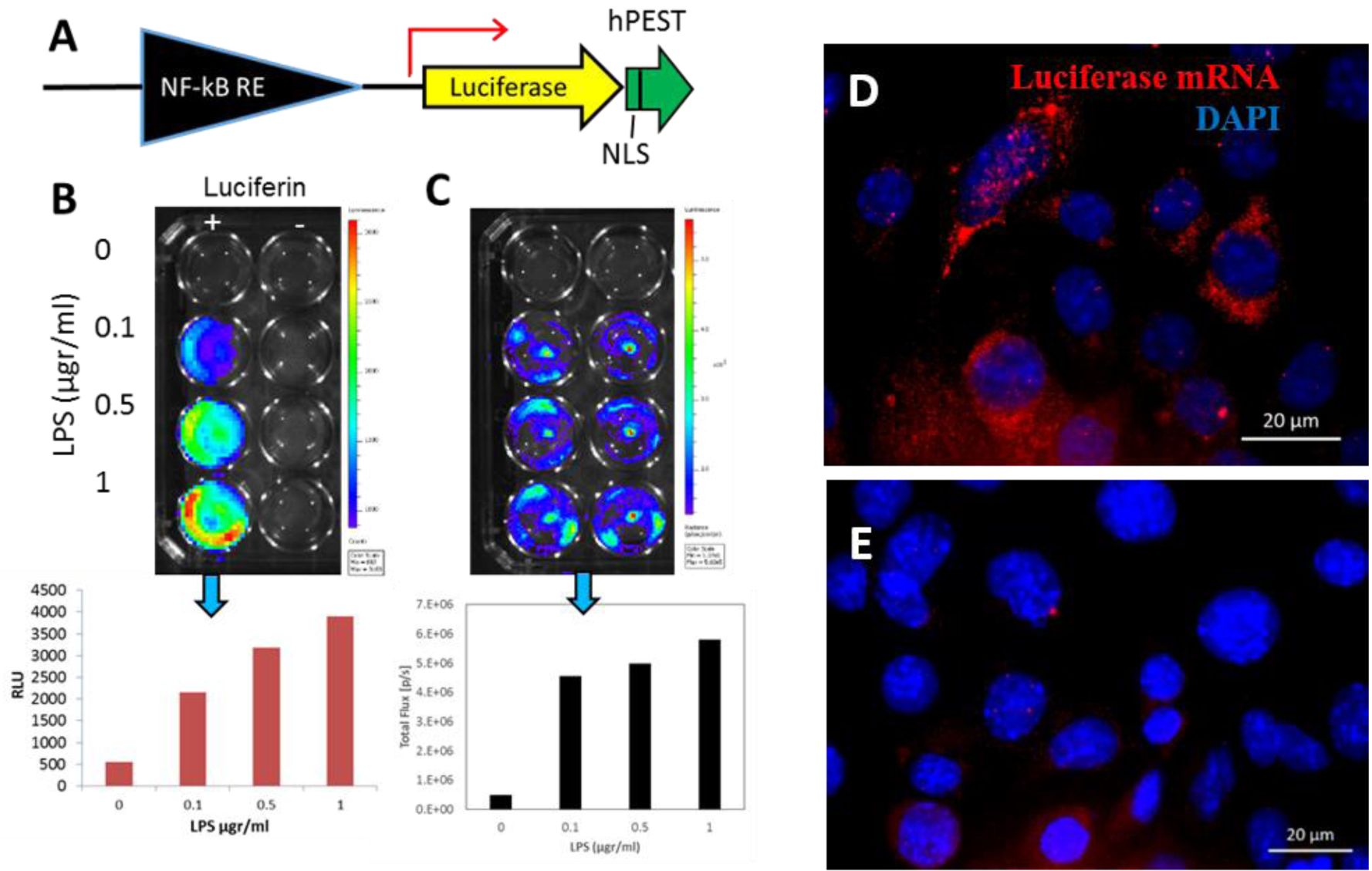
In vitro validation of the lentivirus target and reporter system. (A) demonstrating expression and transcription of luciferase using luminescence analysis (B-C) and fluorescence in situ hybridization (FISH; D-E) of luciferase (luc2CP) mRNA. Human THP-1 (B) and mouse RAW 264.7 (C) macrophage-like cell lines, were infected with the lentiviral NF-κB reporter vector (A). Cells were activated with LPS (B-C), and *Mycoplasma bovis* lipoproteins (D). Luminescence imaging and analysis were performed using cooled CCD optical macroscopic imaging system (B-C; IVIS Lumina Series III, PerkinElmer Inc., MA USA). FISH and nuclear staining with 4,6-diamidino-2-phenylindole (DAPI) of activated (C) and control EPH4 cells (D) demonstrating luc2CP transcripts in activated cells. Microscopic images were taken with epifluorescence microscopy system (Axio Imager M1, Zeiss, Germany). Scale bars 20 µm (C-D).

**Supplementary Figure S2.**
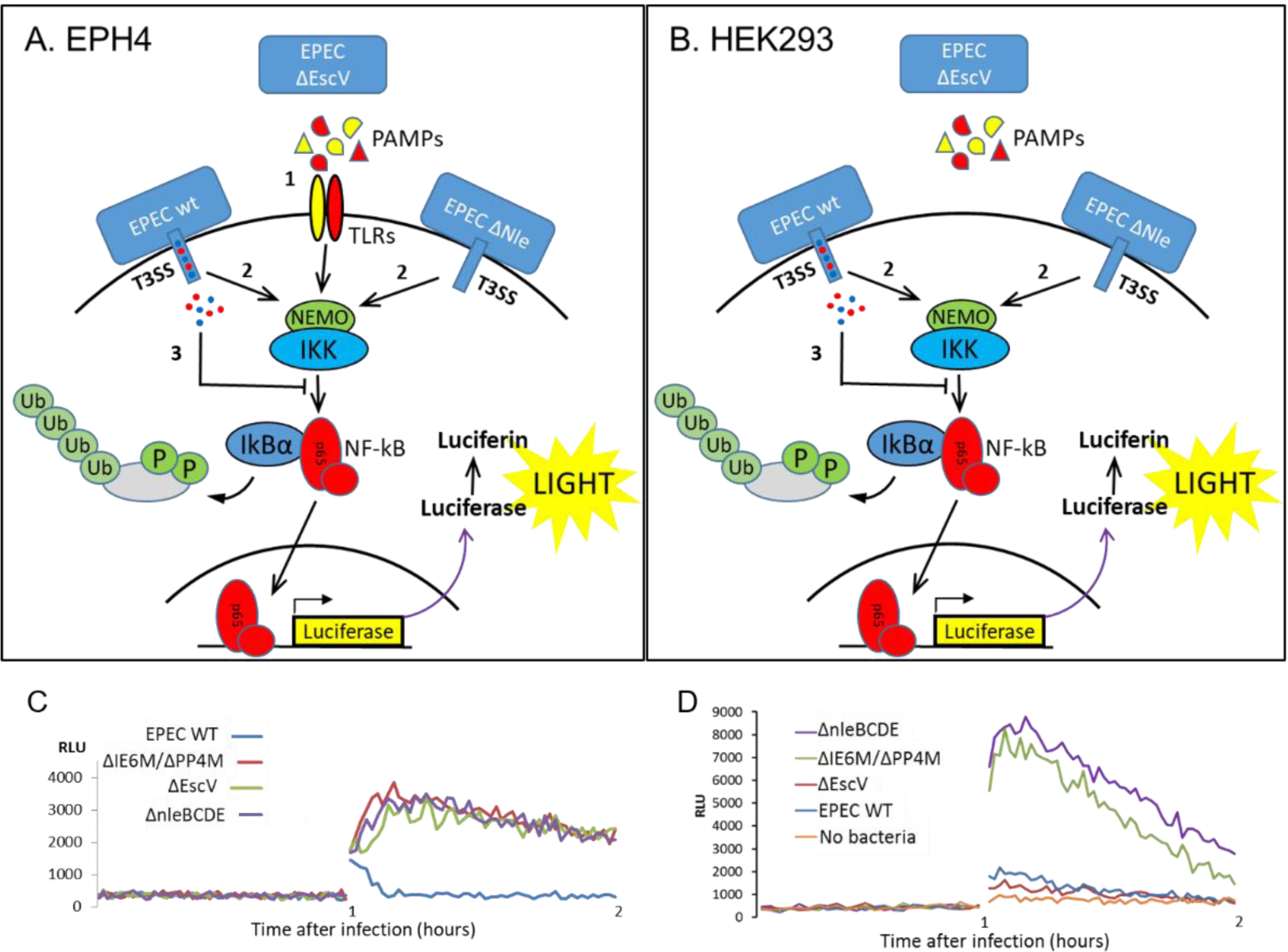
In vitro validation of the lentivirus target and reporter system. Mouse mammary epithelial line EPH4 (A&C) and human embryonic cell line HEK293T (B&D) were infected with the lentiviral NF-κB reporter vector (Figure S1A). Transduced cells were infected with wild type (WT) enteropathogenic *E. coli* bacteria, the type 3 secretion system (T3SS)-defective mutant strain ΔescV, and ΔIE6/PP4, and ΔnleBCDE mutant strains which are defective in anti-NF-κB effectors. In EPH4 cells NF-κB pathway is activated by all bacterial strains through pathogen associated molecular patterns (PAMPs)-toll-like receptor (TLR) signalling (1 in A) and T3SS-dependent activation following infection with WT and the anti-NF-κB-defective mutant strains (2 in A). Inhibition of NF-κB activation is mediated by T3SS-dependent anti-NF-κB effectors and is limited to WT infection of EPH4 cells (3 in A). HEK293T cells are TLR signaling-defective and both activation (2 in B) and inhibition (3 in B) of NF-κB signaling is T3SS-dependent. Luminescence analysis of NF-κB activation following infection with the above described bacterial strains were performed using SpectraMax i3x multiple detection microplate reader (Molecular Devices, CA USA) and are presented in C-D. Representative results of one experiment out of three similar experiments.

**Supplementary Figure S3.**
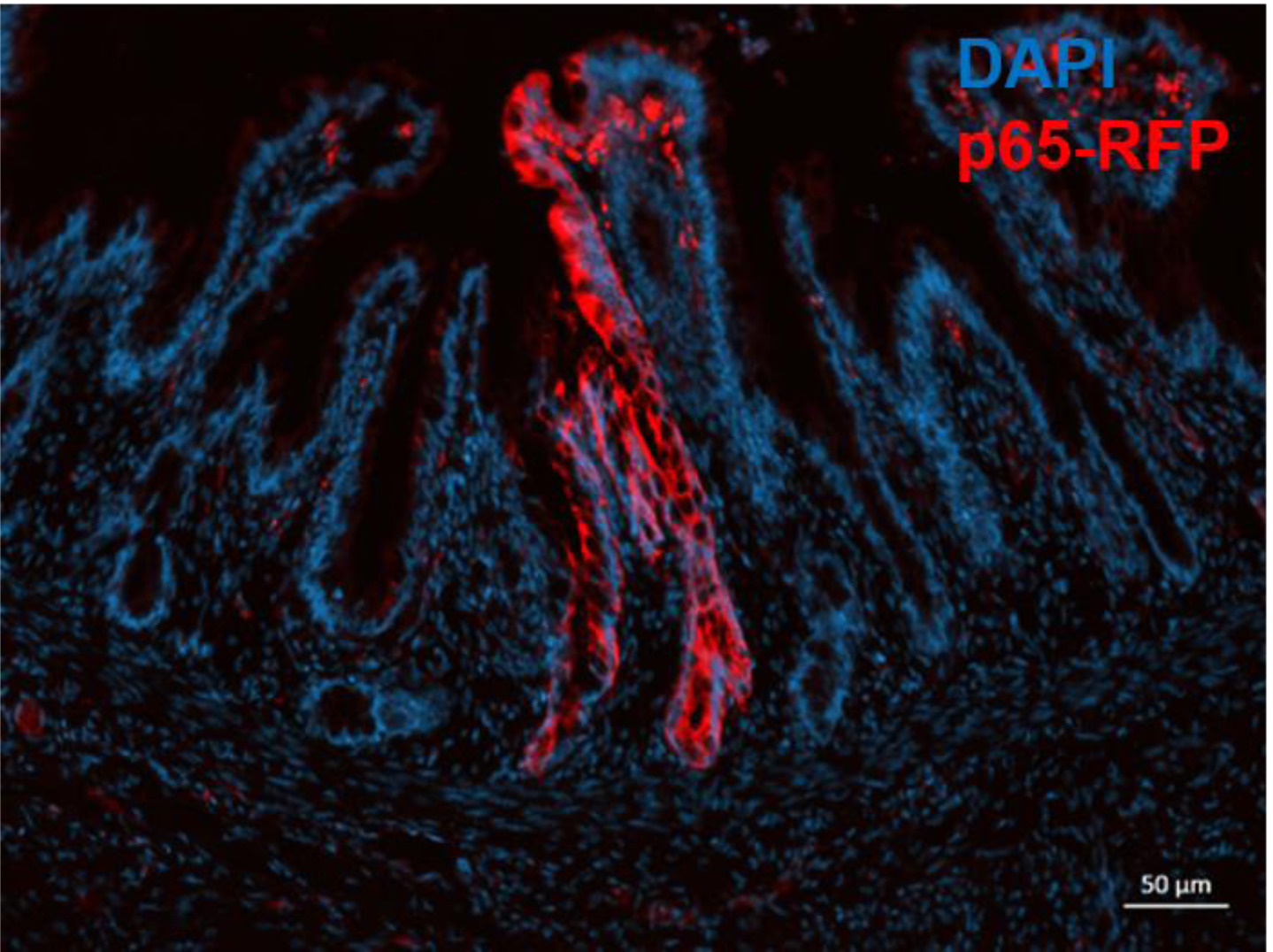
Lineage tracing of human intestinal stem cells (ISC) in gut xenografts. Human fetal gut was infected with lentivirus carrying the human p65-RFP gene. Fetal gut was transplanted subcutaneously in SCID mouse and allowed to develop over 12 weeks into a mature pediatric gut. Cryosections of the gut xenograft were stained with DAPI imaged under fluorescence microscopy. The heritable genetic marker was integrated into stem cells via lentiviral vector infection, all other cells transduced by the lentivirus vector were lost in the course of xenograft development over a 12 week period. The transduced stem cells in which the p65-RFP transgene have been integrated generated progeny that produce the p65-RFP protein. These cells extend from the crypt base to the villus tip and can be visualized as ribbons of red fluorescing cells.

